# Plausible pathway for a host–parasite molecular replication network to increase its complexity through Darwinian evolution

**DOI:** 10.1101/2022.01.17.476531

**Authors:** Rikuto Kamiura, Ryo Mizuuchi, Norikazu Ichihashi

## Abstract

How the complexity of primitive self-replication molecules develops through Darwinian evolution remains a mystery with regards to the origin of life. Theoretical studies have proposed that coevolution with parasitic replicators increases network complexity by inducing inter-dependent replication. However, the feasibility of such complexification with biologically relevant molecules is still unknown owing to the lack of an experimental model. Here, we investigated the plausible complexification pathway of host–parasite replicators using both an experimental host–parasite RNA replication system and a theoretical model based on the experimental system. We first analyzed the parameter space that allows for sustainable replication in various replication networks ranging from a single molecule to three-member networks using computer simulation. The analysis shows that the most plausible complexification pathway from a single host replicator is the addition of a parasitic replicator, followed by the addition of a new host replicator that is resistant to the parasite. We also provide evidence that the pathway actually occurred in our previous evolutionary experiment. These results provide both a theoretical basis and experimental evidence that a single replicator spontaneously develops into multi-replicator networks through coevolution with parasitic replicators, which might be the first complexification step toward to the emergence of life.

## Introduction

Most of the origin of life scenarios hypothesize that a simple self-replicating molecule or a set of replicating molecules appeared and underwent Darwinian evolution to gradually become more complex toward the extant life (1–5). To examine the plausibility of this scenario, researchers have synthesized self-replication molecules such as simple RNA or peptides, which might have been available on the early Earth (reviewed in 4–6), although Darwinian evolution of these simple molecules remains a challenge. RNA or DNA replication systems capable of Darwinian evolution have been constructed using proteins of the existing organisms (8–11). Although the proteins used in these systems did not exist on the early Earth, they could be utilized as experimental models that might mimic some aspects of primitive replicators consisting of biologically relevant molecules such as RNA and peptides. Even for replication systems consisting of modern proteins, however, the development of complexity through Darwinian evolution, a prerequisite for the emergence of life, remains a significant challenge.

Complexity is an ambiguous concept, and there are several measures for determining the complexity of a replication system, such as the amount of information encoded in a replicator (12), the number of traits of a replicator (13), the difficulty in achieving traits (14), and the number of replicators organized as a replication network (15,16). Here, we focus on one of the measures, the number of replicators in a replication network (i.e., network complexity). One of the possible pathways for a replicator to develop this complexity is the diversification of replicators and formation of inter-dependent, cooperative replication networks among them, such as a hypercycle (2,17–21). A major hurdle for inter-dependent network formation is parasitic replicators, which destroy the cooperative replication network (17,22). To date, theoretical (23–27) and experimental (28,29) studies have revealed that spatial structures such as compartmentalization repress parasitic replicators. Furthermore, recent theoretical studies conducted by Takeuchi and Hogeweg showed that parasitic replicators induced diversification of RNA-like replicators through evolutionary arms race in compartmentalized structures and allowed the formation of more complex inter-dependent molecular networks (15). These studies suggest that parasitic replicators, which have been considered as an obstacle to complexification, may play an important role in complexification in compartmentalized structures. One of the remaining challenges is the plausibility of such complexification within a realistic parameter space that is achievable with biologically relevant molecules such as RNA and proteins.

Recently, we constructed an in vitro translation-coupled RNA replication system and demonstrated the coevolution of host and parasitic RNAs (29,30). In this system, a host RNA replicates through the translation of the self-encoded replication enzyme, whereas parasitic RNAs, which spontaneously appear, replicate by relying on the replication enzyme translated from the host RNAs. When the replication was repeated through serial replication processes in water-in-oil compartments, the RNAs were mutated by replication errors and underwent Darwinian evolution. In a previous study, we showed that host and parasitic RNAs diversified into multiple lineages through Darwinian evolution (30). In a recent study, we further repeated the serial replication processes and found that the diversified RNA species start to co-replicate by forming an inter-dependent network, which finally consists of three hosts and two parasites (31). These experimental results support the idea that coevolution between host and parasitic replicators can drive diversification and complexification. However, it is still unknown how such complexification is possible and competitive exclusion among RNA species is circumvented.

This study addressed two questions. 1. How do multiple hosts and parasites sustainably co-replicate by avoiding competitive exclusion? 2. What is the plausible pathway for complexification? To this end, we first constructed a theoretical model of a compartmentalized host–parasite replication system based on our experimental method and parameters. We then investigated the parameter space that allows for the sustainable replication of all replicators in each host–parasite replication network in up to three-member networks using computer simulation. We also conducted an evolutionary simulation by introducing new replicators with different parameters. These simulations showed that the parasites mediate the co-replication of multiple host species as a “niche.” The simulations also showed that the most plausible complexification pathway from a single host is the successive addition of a parasite first and then a new host that is resistant to the parasite, although the other pathways are also possible in narrower ranges of parameter space. Furthermore, we confirmed that the most plausible pathway (the addition of a parasite, followed by the addition of a parasite-resistant host) occurred in our previous evolutionary experiment.

## Results

### Strategy of theoretical model and analysis

Fig. 1A shows a possible complexification pathway for host–parasite replication networks with up to three members. A single host replicator, termed “H,” possibly forms two-member replication networks by the addition of a new host or parasite, termed HH and HP networks, respectively. The next step is to form the three-member networks, named HHH, HHP, and HPP. In this study, we evaluated the feasibility of each network by comparing the parameter spaces that allow for the sustainable replication of all members in the networks for certain generations. The parameters used here are replication coefficients, in which each host replicates itself or other replicators. For example, the HP network can be characterized by two replication coefficients, that for self-replication (k_H_) and that for the replication of the parasite (k_P_) (Fig. 1B).

**Figure 1.**
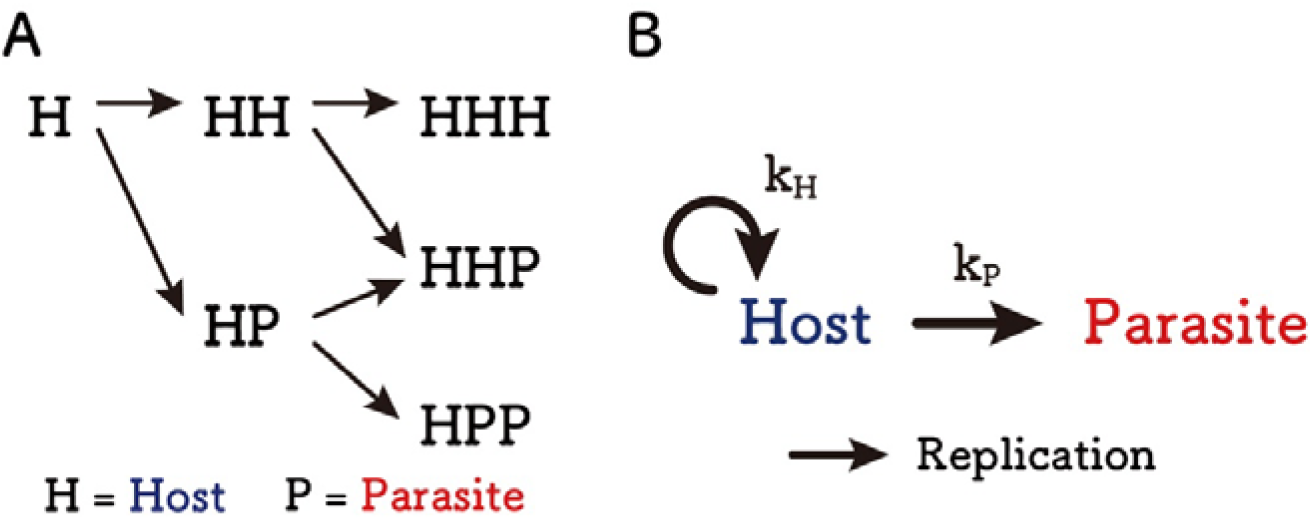
Possible complexification pathways and an example of replication parameters. (A) Possible complexification pathways in up to three-member replication networks. Starting from a single host self-replicator (H), next possible steps are the addition of another host or parasite to form two-member replication networks, namely, HH and HP. In the next step, another host or parasite could join to form three-member replication networks, namely, HHH, HHP, and HPP. (B) Parameters that characterize HP network as an example. A host replicates itself (i.e., self-replicates) with the coefficient k_H_ and a parasite with the coefficient k_P_. Similar coefficients are used for the other networks.

To investigate the parameter space that allows for the sustainable replication of all members in a replication network, we constructed a theoretical model of compartmentalized host–parasite replication. The replication is continued by repeating three steps: replication, selection, and fusion-division (Fig. 2). The detailed procedures are described in the Methods section.

**Figure 2.**
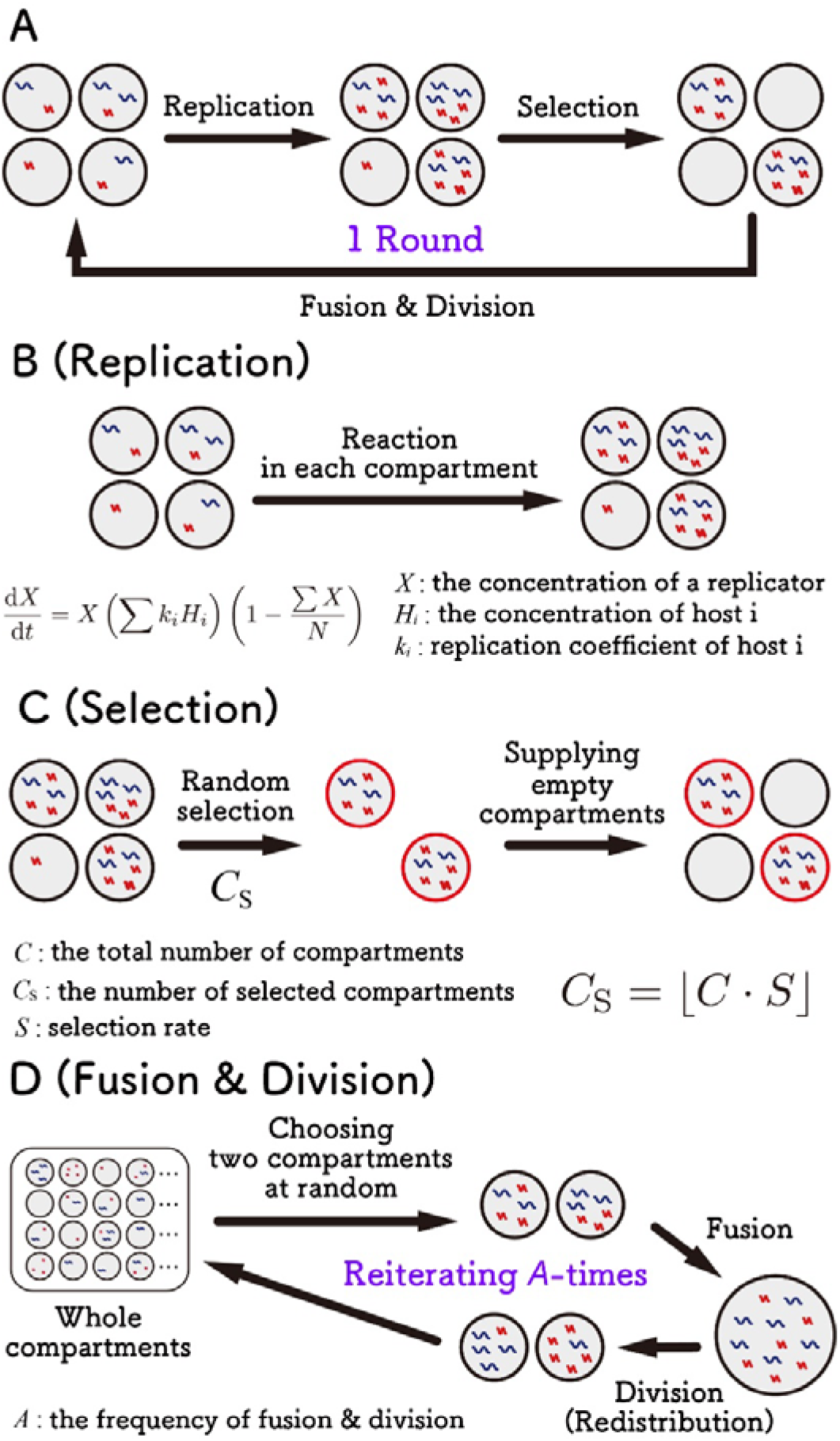
Theoretical model of compartmentalized replication through serial replication cycles. (A) Overview of the serial replication cycle of compartmentalized replication, which consists of replication, selection, and fusion-division steps. (B) In the replication step, hosts and parasites in each compartment replicate according to different equations. (C) In the selection step, a certain number (Cs) of compartments are randomly selected, and the other compartments are replaced with empty compartments. (D) In the fusion-division step, two compartments are randomly chosen, and the internal host and parasites are mixed, followed by random redistribution into two compartments. These processes were repeated *A* times. The number of compartments is 3,000, and the frequency of fusion-division is 5,000 unless indicated otherwise.

The model constructed in this study, which mimics our previous evolutionary experiments (29), differs from those of previous studies in the literature at some points. For example, some replicator models assume compartments aligned in 2D space, where each compartment interacts with only adjacent compartments (15,32), but our compartments are well-mixed and can fuse with any compartment. In other studies, the cellular compartments are assumed to grow and divide depending on the internal reaction (23,24,33), whereas in our model, the volumes of the compartments and the fusion-division steps are independent of the internal reaction.

### HH networks

First, we investigated the HH network, in which two host species (Host 1 and Host 2) self-replicate with coefficients k_11_ and k_22_ and cross-replicate with coefficients k_12_ and k_21_, respectively (Fig. 3A). We performed computer simulations of the compartmentalized replication shown in Fig. 2 with all combinations of four values (1,7, 2.0, 2.3, and 2.6) for each coefficient. The four values are based on experimental data that were obtained in the later experiment conducted in this study (Table 1), in which the maximum and minimum coefficients were approximately 2.3 and 2.0, respectively, and additionally we chose a larger value (2.6) and a smaller value (1.7). We also tested more extreme values (0.2 and 4.1) in Fig. S1.

**Table 1.** Estimated replication coefficients.

|  | Replicating RNA |  |  |  |
| --- | --- | --- | --- | --- |
| | | $\text{Host}^1_{\text{exp}}$ | $\text{Host}^2_{\text{exp}}$ | $\text{Parasite}_{\text{exp}}$ |
| Replicated RNA | $\text{Host}^1_{\text{exp}}$ | 2.3 ($k_{11}$) | 2.3 ($k_{21}$) | 0.0 |
| | $\text{Host}^2_{\text{exp}}$ | 2.3 ($k_{12}$) | 2.0 ($k_{22}$) | 0.0 |
| | $\text{Parasite}_{\text{exp}}$ | 6.7 ($k_{1p}$) | 0.0 ($k_{2p}$) | 0.0 |

\*Corresponding parameter names in the theoretical model are shown in parentheses.

**Figure 3.**
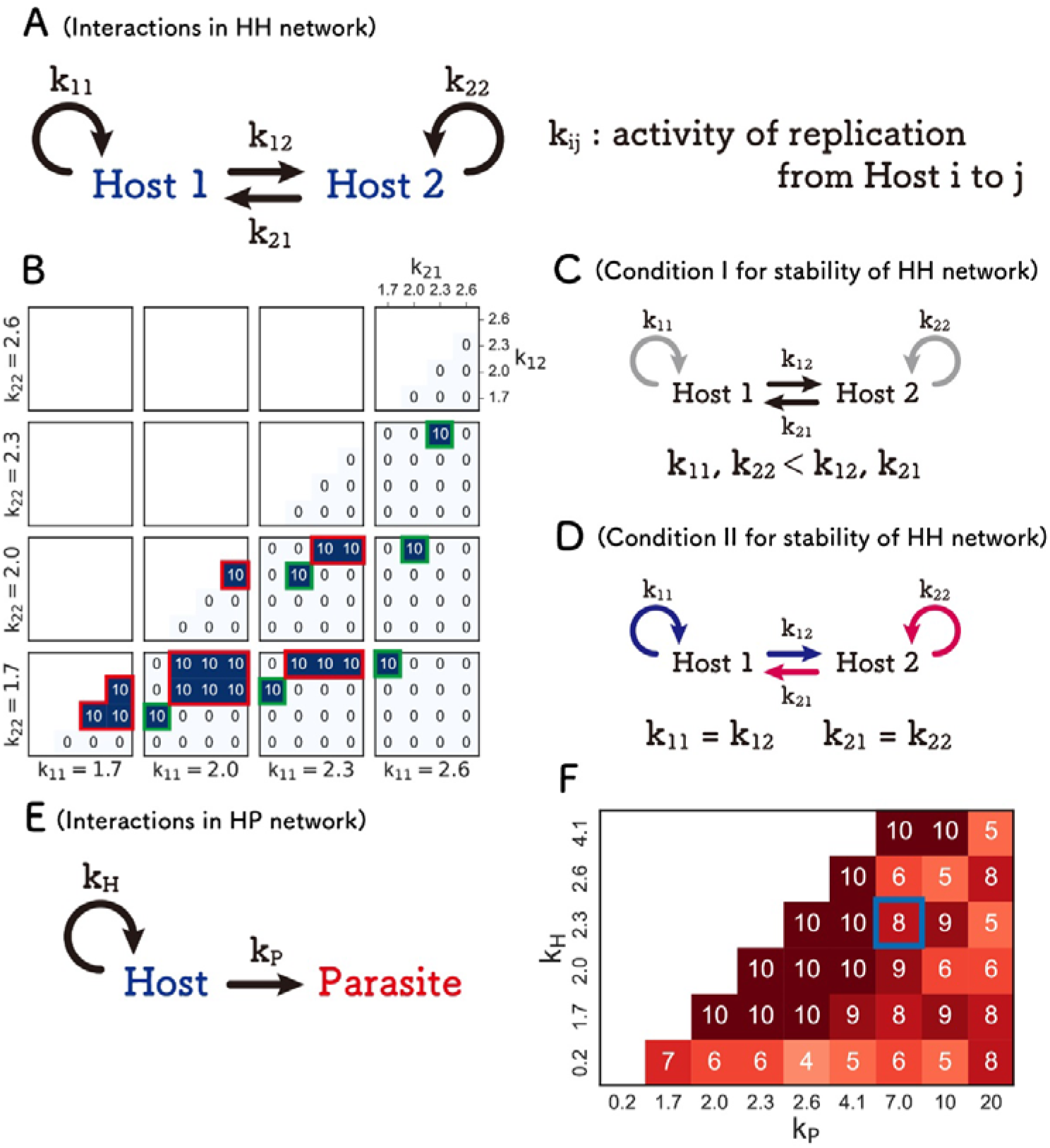
Search for the parameters that allow sustainable HH and HP networks. (A) Scheme of the HH network. Each host self-replicates with coefficient k_11_ or k_22_, and replicates the other host with coefficients k_12_ or k_21_. (B) Numbers of the runs in which both Hosts 1 and 2 are sustained for 100 rounds out of 10 independent simulations. The regions enclosed with red and green squares are the two different sustainable conditions each depicted in (C) and (D), respectively. The results on the diagonal line were omitted because Hosts 1 and 2 are identical there. (E) Scheme of the HP network. The host self-replicates with coefficient, k_H_, and replicate the parasite with coefficient, k_P_. (F) Numbers of the runs in which both the host and parasite are sustained for 100 rounds out of 10 independent simulations. The blue square indicates parameters of the dominant RNAs (Host1_exp_ and Parasite_exp_) obtained in the evolutionary experiment.

Using the combinations of these coefficients, we performed 10 independent simulations and counted the number of “sustained” runs in which all replicators were sustained for 100 rounds of the serial replication cycle (Fig. 3B). With most of the parameter sets, the HH replication network was not sustained even once, suggesting that the co-replication of two types of host species is unlikely. However, there are some parameter sets that allow sustained replications for all 10 runs, which can be categorized into two conditions. Condition I is a “low self- and high cross-replications” condition (i.e., *k*_12_, *k*_21_ > *k*_11_, *k*_22_, Fig. 3C), where the two hosts replicate cooperatively, marked with red squares in Fig. 3B. Condition II is a “balanced replication” condition (*k*_11_ = *k*_12_, *k*_21_ = *k*_22_, Fig. 3D, where one host species self-replicates as much as cross-replicates with the other host species, marked with green squares. These results indicate that sustainable replication in the HH network occurs only in limited cases. The results are similar for the extreme parameters (Fig. S1).

### HP networks

Next, we investigated the HP network, in which a host self-replicates with coefficients k_H_ and replicates a parasite with coefficients k_P_ (Fig. 3E). For these replication coefficients, we used the same values as those used in the HH network, including the two extreme values (0.2, 1,7, 2.0, 2.3, 2.6, and 4.1). Additionally, we adopted the experimental value (7.0) and larger values (10.0 and 20.0) for the coefficients of the parasite (k_P_). Using the combinations of these coefficients, we performed a series of computer simulations and counted the number of sustained runs out of 10 independent runs (Fig. 3F). The number of sustained runs was at least four in all parameter sets tested here, and gradually changed within the parameter space. This result exhibits a clear contrast to the HH network in Fig. 3B, in which the sustained parameter was relatively rare, and the number of sustained runs changed sharply with a small parameter change. These results indicate that the HP network is more sustainable in a broader parameter space than the HH network.

The number of sustained runs may depend on the number of compartments. To test this hypothesis, we conducted simulations with a varied number of compartments, *C*. The frequency of fusion-division, *A*, was determined so that the frequency of fusion-division per compartment was a fixed value. In the HP network (Fig. S2A), the number of sustainable replications increased when the number of compartments was increased from 3,000 to 10,000 (the original number was 3,000). This is probably because a large number of compartments provide hosts with a greater chance of escaping from parasites. Similarly, in the HH network (Fig. S2B), the parameter sets that allow sustainable replication slightly increased when the number of compartments was increased from 3000 to 10,000 (compare Fig. 3B with Fig. S2B left and Fig. S1 with Fig. S2B right), while the two hosts were still unsustainable, with most of the parameter sets. These results confirm that the HP network is sustainable in a broader parameter space than the HH network, even with a larger number of compartments.

### HPP network

Next, we investigated the HPP network, in which another parasite was added to the HP network, with the same simulation method and parameters as for the HP network (Fig. 4A). The number of sustained runs, in which all host and parasites are sustainably replicated for 100 rounds, in 10 independent simulations, are shown in Fig. 4B. The host and two parasites are never sustainably replicated together with any combinations of parameters used here, mainly due to the competition between the two parasites, which immediately exterminates one with a smaller coefficient. Even after increasing the number of compartments to 10,000, no sustainable replication was observed (Fig. S2C). These results suggest that even if an HPP network is formed during evolution, it will soon return to the HP network.

**Figure 4.**
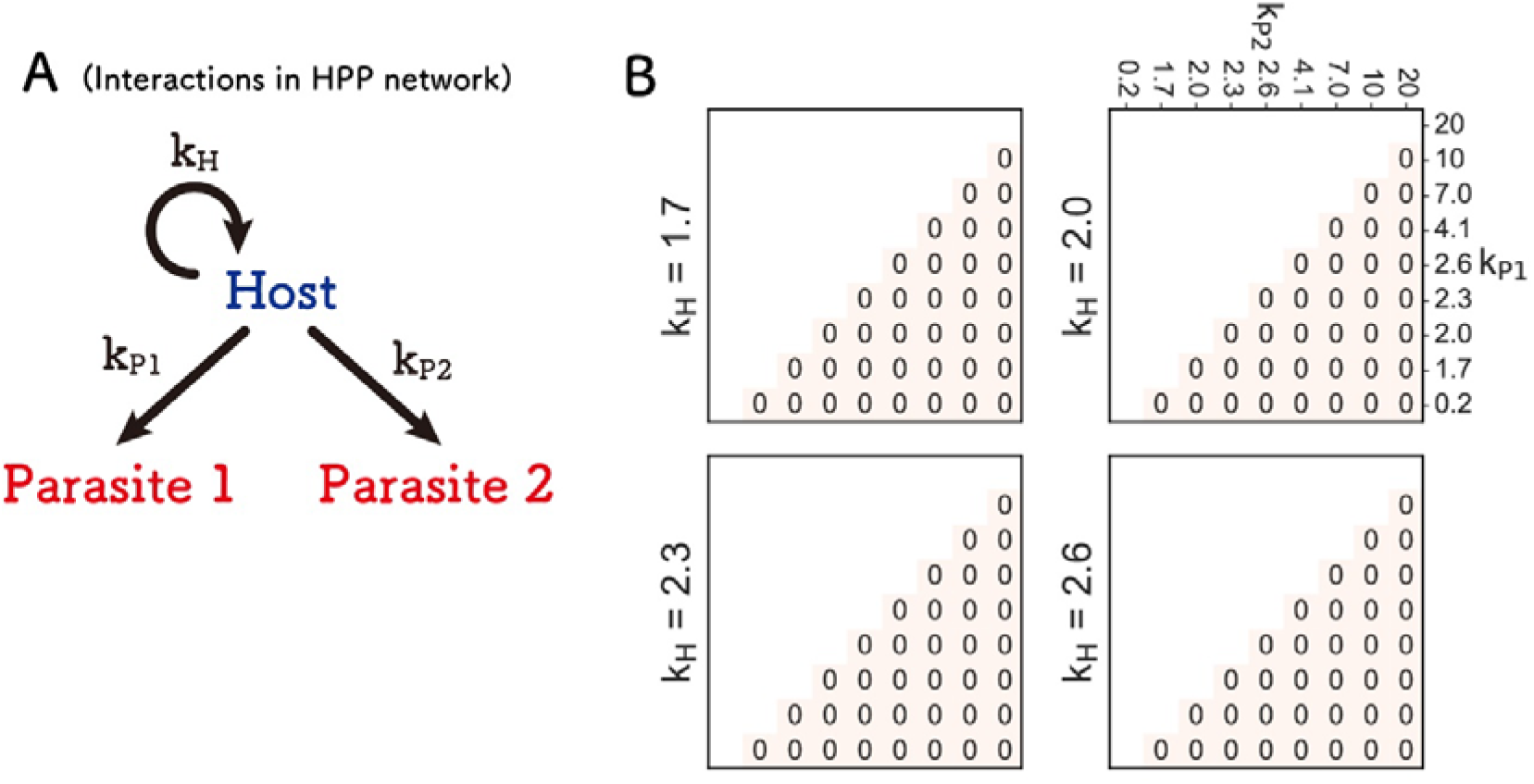
Search for the parameters that allow sustainable HPP network. (A) Scheme of the HPP network. The host self-replicates with coefficient kH and replicates two parasites with coefficient k_P1_ or k_P2_. (B) Numbers of the runs in which all three replicators (the host and Parasites 1 and 2) are sustained for 100 rounds out of 10 independent simulations.

### HHP network

Next, we investigated the HHP network, in which another host was added to the HP network or another parasite was added to the HH network, with the same parameter range as in the HH network. Since there were too many parameter combinations to compute in a reasonable time, we used only two types of parasite coefficients: the same coefficient for both hosts (k_1P_ and k_2P_ = 7.0, Fig. 5A), termed “symmetrical parasite replication,” where both hosts replicate the parasite similarly, or much smaller coefficient for one of the hosts (k1P = 7.0 and k2P = 0.1, Fig. 5C), termed “asymmetrical parasite replication,” where one of the hosts is resistant to the parasite. The value (7.0) was adopted from the experimental data, and the value (0.1) was chosen as an example of much smaller values. We also simulated the case with an intermediate value of 1.0, but the results were similar (Fig. S3).

**Figure 5.**
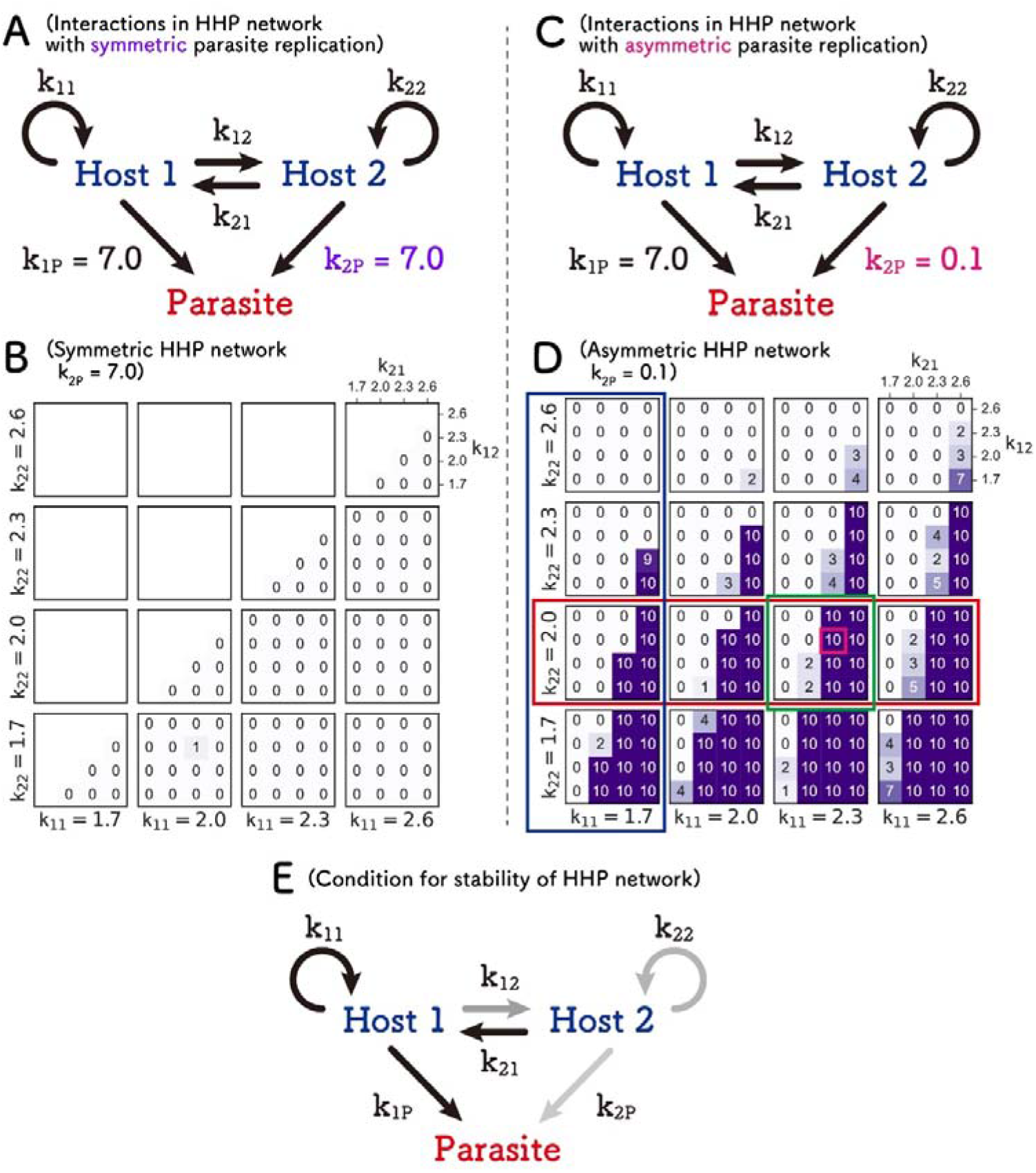
Search for the parameters that allow sustainable HHP network. Symmetric (A) and asymmetric (C) HHP networks The number of runs in which all three replicators (Hosts 1 and 2, and the parasite) were sustained for 100 rounds out of 10 independent simulations in symmetric (B) and asymmetric (D) cases. The magenta square represents the close parameter values of the representative RNAs obtained from the evolutionary experiment. (E) A typical condition for a sustainable HHP network, which contains a parasite-susceptible and parasite-resistant host species, and the parasite-susceptible host (Host 1) tends to replicate more efficiently through self- and/or cross-replications. The color depth of the arrows represents the value of the replication coefficients.

The number of sustainable replications out of 10 independent runs was strikingly different between the symmetrical and asymmetrical cases; in the symmetrical case (Fig. 5B), sustainable replication was rarely observed with the parameter sets we tested. This is due to the competition between the two hosts; when the two hosts have the same susceptibility to the parasite, a host that replicates more competitively excluded the other host. By contrast, there were many parameter sets that allows sustainable replications, termed “sustainable parameters,” in the asymmetrical case (Fig. 5D). We also obtained a similar result with the extreme parameters (Fig. S4). These results indicate that the HHP network can be sustainable with a certain range of parameters when parasite resistance is asymmetrical between the two hosts. Furthermore, it should be noted that the sustainable parameters in the HHP network overlap with those in the HP network (i.e., the sustainable Host 1 and parasite in the HHP network are also sustainable in the HP network) because we used the same parameter range, which implies that a sustainable HP network can form a sustainable HHP network soon after the appearance of a parasite-resistant host. These results suggest that the transition from HP to HHP networks is a plausible pathway for complexification in the replication network.

We examined the asymmetric case in more detail. First, we found that as the self-replication of Host 1 (non-resistant host) increased (i.e., *k*_11_ increased), the number of sustainable runs gradually increased. For example, in the parameter region of *k*_22_ = 2.0 (red rectangle in Fig. 5D), the region with 10 points increased from left to right (in the direction of increasing *k*_11_). By contrast, as the self-replication of Host 2 (resistant host) increased (i.e., *k*_22_ increased), the number of sustainable runs decreased. For example, in the region of *k*_11_ = 1.7 (blue rectangle), the region with 10 points significantly decreased from bottom to top (in the direction of increasing *k*_22_). Second, we found that as the cross-replication of Host 2 to Host 1 (*k*_21_) increased, the number of sustainable runs increased. For example, in the region of *k*_11_ = 2.3 and *k*_22_ = 2.0 (green rectangle), the region with 10 points increases from left to right (in the direction of increasing *k*_21_). In summary, a sustainable asymmetric HHP network requires parameter sets that favor replication of the parasite-susceptible host either by self-replication or cross-replication (a typical condition is schematically depicted in Fig. 5E).

### HHH network

We investigated the HHH network (Fig. 6A). Because there are too many parameters to simulate in realistic time in this network, we fixed the parameter values for Hosts 1 and 2 (k_11_, k_21_, k_12_, and k_22_) in two cases that allow sustainable replication in the HH network (Fig. 3B): one of the Conditions I (i.e., “low self- and high cross-replications” conditions) (k_11_ = 1.7, k_21_ = 2.6, k_12_ = 2.0, and k_22_ = 1.7) and one of the Conditions II (i.e., “balanced replications” conditions) (k_11_ = 2.0, k_21_ = 1.7, k_12_ = 2.0, and k_22_ = 1.7). In these two cases, we searched for sustainable parameter sets for the new Host 3. All combinations of four values (1.7, 2.0, 2.3, and 2.6) were tested for the four new replication coefficients (k_31_, k_32_, k_13_, and k_23_) generated by the addition of Host 3. Only the smallest and largest values (1.7 and 2.6) were used for k_33_, the self-replication coefficient of Host 3. The number of independent runs was reduced from ten to three to decrease the computational cost.

**Figure 6.**
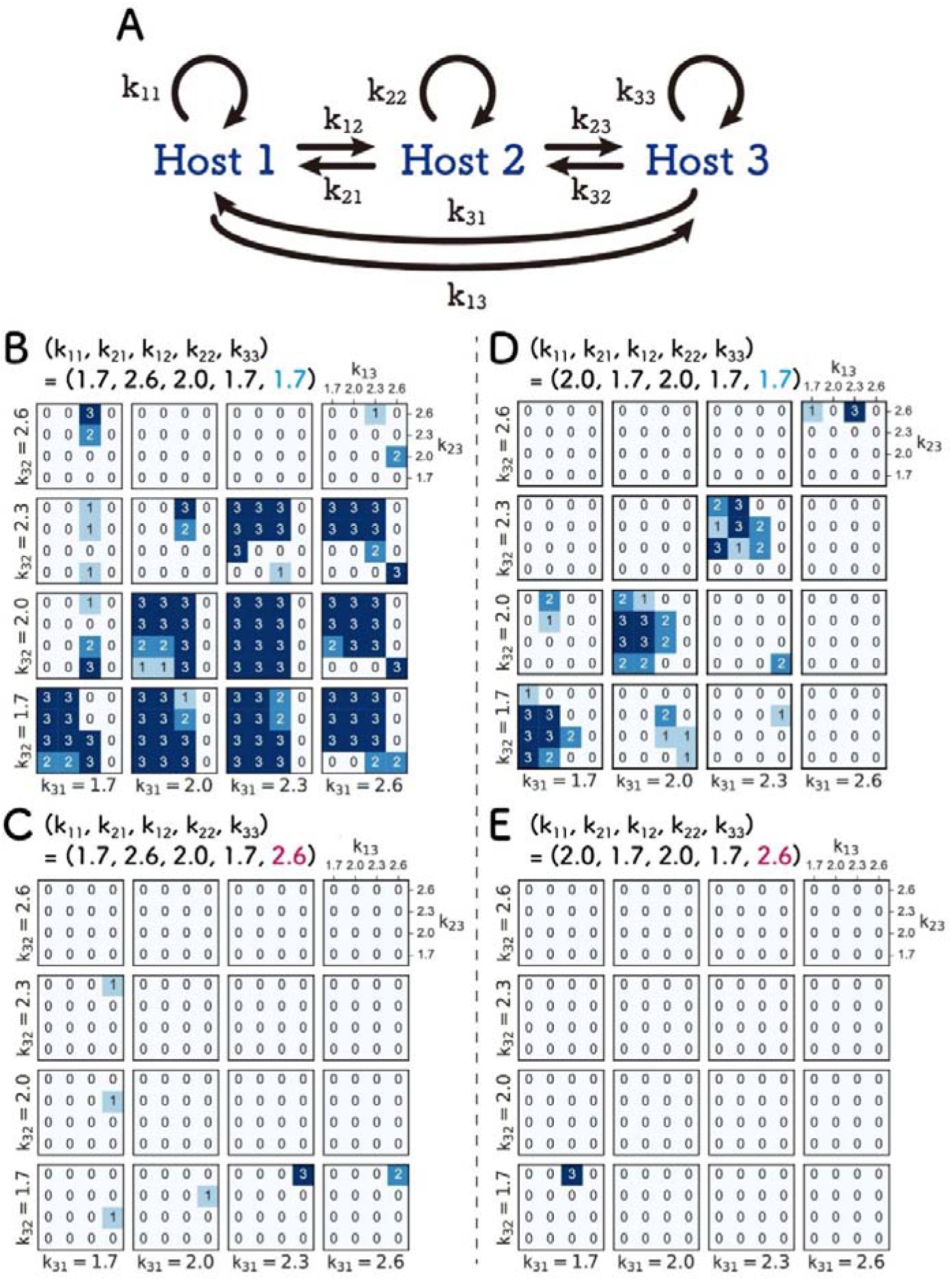
Search for the parameters that allow sustainable HHH network. (A) Scheme of the HHH networks. (B-E) Numbers of runs in which all three hosts are sustained for 100 rounds out of three independent simulations. The parameter values for Hosts 1 and 2 are fixed at two cases that allows sustainable replication in the HH network. (B, C) Conditions I (Fig. 3C, k_11_ = 1.7, k_21_ = 2.6, k_12_ = 2.0, and k_22_ = 1.7). (D, E) Conditions II (Fig. 3D, k_11_ = 2.0, k_21_ = 1.7, k_12_ = 2.0, and k_22_ = 1.7). For the same reason, we employed a small (1.7) (B and D) or a large (2.6) (C and E) value for the self-replication coefficient of the newly added Host 3 (k_33_).

Under the “low self- and high cross-replications” conditions (Figs. 6B and 6C), sustainable parameter sets were frequently found when k_33_ is the smaller value, 1.7 (Fig. 6B), whereas barely found when k_33_ is the larger value, 2.6 (Fig. 6C), indicating that low self-replication also for the new Host 3, which induces inter-dependent replication of all replicators, is important for the sustainability.

A similar trend was also found under the “balanced replication” conditions (Figs. 6D and 6E), the three hosts sustainably replicated with a certain range of parameter space with the lower k_33_ values (Fig. 6D), whereas such parameter sets were rarely found with the larger k_33_ values (Fig. 6E). We found that when the parameters for Host 3 are also “balanced” (i.e., k_11_ = k_12_ ≈ k_13_, k_22_ = k_21_ ≈ k_23_, and k_33_ ≈ k_31_ = k32), the HHH network tended to be sustainable under these conditions.

In summary, when we added another host to the sustainable HH network, the resultant HHH network could be sustainable again with certain parameter sets, implying that host-only networks are plausible even when members of the network increase. However, a large obstacle for the formation of HH and HHH networks is the appearance of parasitic replicators, which is inevitable, at least in our experimental model. This point is further discussed in the discussion section.

### Computer simulation of the transition of the networks

Next, we investigated the possible evolutionary transition of the networks using computer simulation. We introduced a mutation step in the serial replication cycle immediately before the replication step, as shown in Fig. 2. In the mutation step, a new host or parasite appears at a certain rate in one of the compartments if the total number of replicator species in the system is less than three. A new replicator is a host or a parasite at the ratio of 0.2 or 0.8, respectively. The new replicator has parameter sets randomly chosen from certain values within the parameter ranges in the simulation conducted above. Starting from a single host species (k_11_ = 2.0), we performed 3000 rounds of serial replication cycles for 100 times and found that in most of the runs (97 runs), all replicators were diluted out, while in three runs, the HHP network was formed. The trajectories of the replicator concentrations for one of the three cases are shown in Fig. 7, in which the HP network first formed between a host (Host 1) and a parasite (Parasite) around the round 400, and then a new parasite-resistant host (Host 2) joined to form the HHP network. These results indicate that sustainable replication networks are rarely formed spontaneously, at least with the parameter sets and the number of compartments used here; however, when they are formed, the formation of an HP network followed by an HHP network is a favorable pathway.

**Figure 7.**
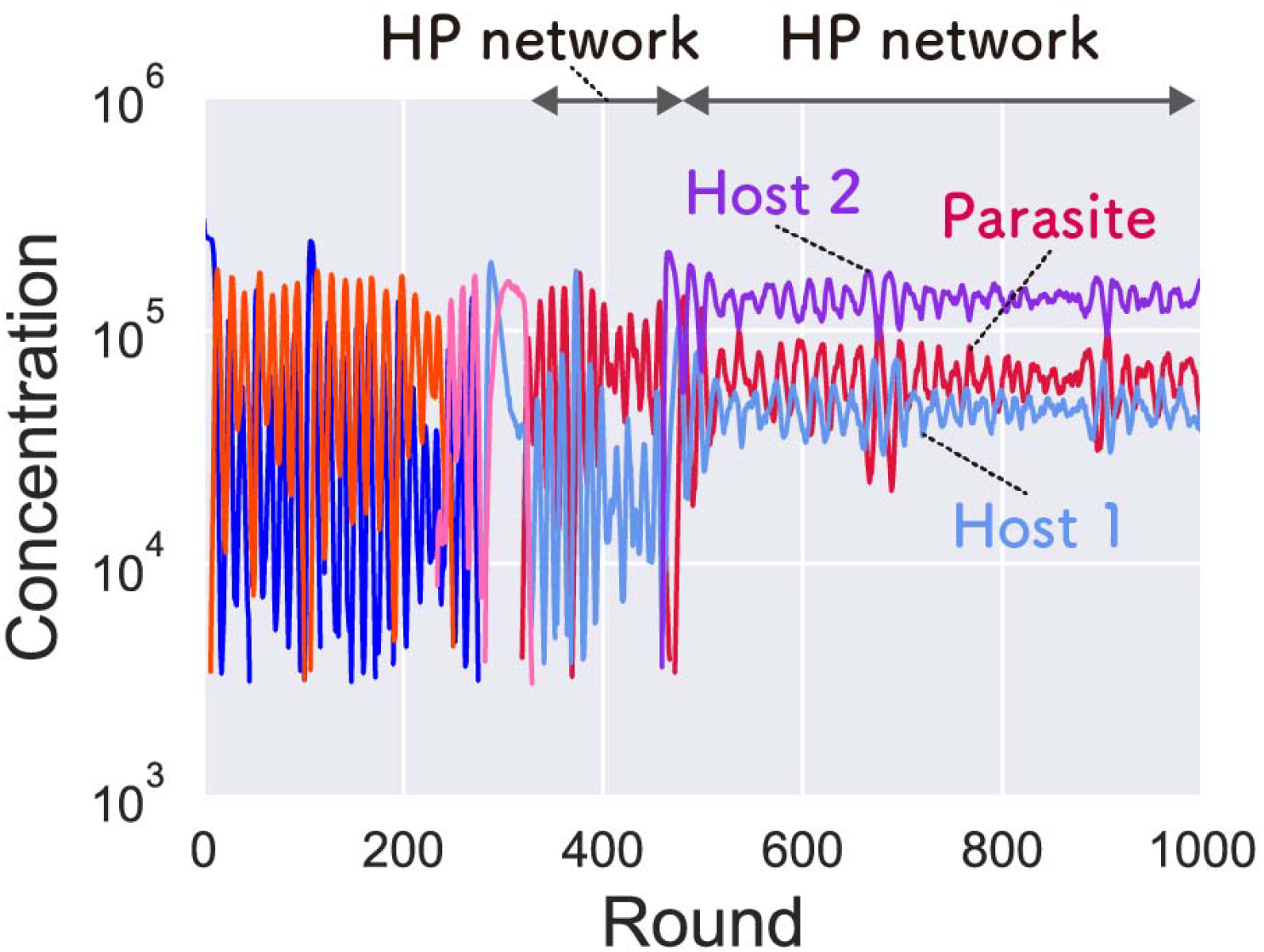
Computer simulation of the evolutionary transition of replication networks. Evolutionary transition was simulated by introducing a mutagenesis step in the serial replication cycle, as shown in Fig. 2. The HHP network was formed in three out of 100 simulations. The trajectory of the concentrations in one of the three runs is shown. Concentrations of more than 3000 were plotted. The parameters of Host 1, Host 2, and parasite were as follows: k_11_ = 2.6, k_12_ = 2.4, k_1P_ = 8.0, k_21_ = 2.5, k_22_ = 2.0, and k_2P_ = 0.1. The HHP networks were sustained for at least 3000 rounds.

### Host and parasitic RNAs that may form HP and HHP networks in the previous evolutionary experiment

In our previous serial replication experiments of compartmentalized translation-coupled RNA replication, we found that a parasitic RNA appeared soon after starting replication and co-replicated with the original host RNA (29). During further serial replication cycles, the host RNA diversified into two distinct lineages (30). These results suggest that sustainable HP and HHP networks might have been formed during the evolutionary experiment, which is consistent with the simulation results. To confirm this possibility, we tested whether the dominant host and parasitic RNAs that appeared during the experiment had replication parameters that support sustainable HP and HHP networks.

To isolate dominant RNAs that possibly form HP and HHP networks, we analyzed the sequence data obtained in a previous study (30). We focused on the early period (up to 39 rounds), where the host RNA starts to diversify. We chose the top eight most frequent sequences of each of the RNA populations in these rounds and drew phylogenetic trees for both the host and parasitic RNAs (Figs. 8A and 8B), along with heat maps that represent the frequencies of each sequence (Figs. 8C and 8D). For parasitic RNA, the phylogenetic tree (Fig. 8A) and the frequency (Fig. 8C) did not show any clear trends, but the most dominant parasitic RNA at round 13 remained as one of the dominant sequences until round 33. We chose this RNA (indicated as Parasite_exp_) as representative of the parasite.

**Figure 8.**
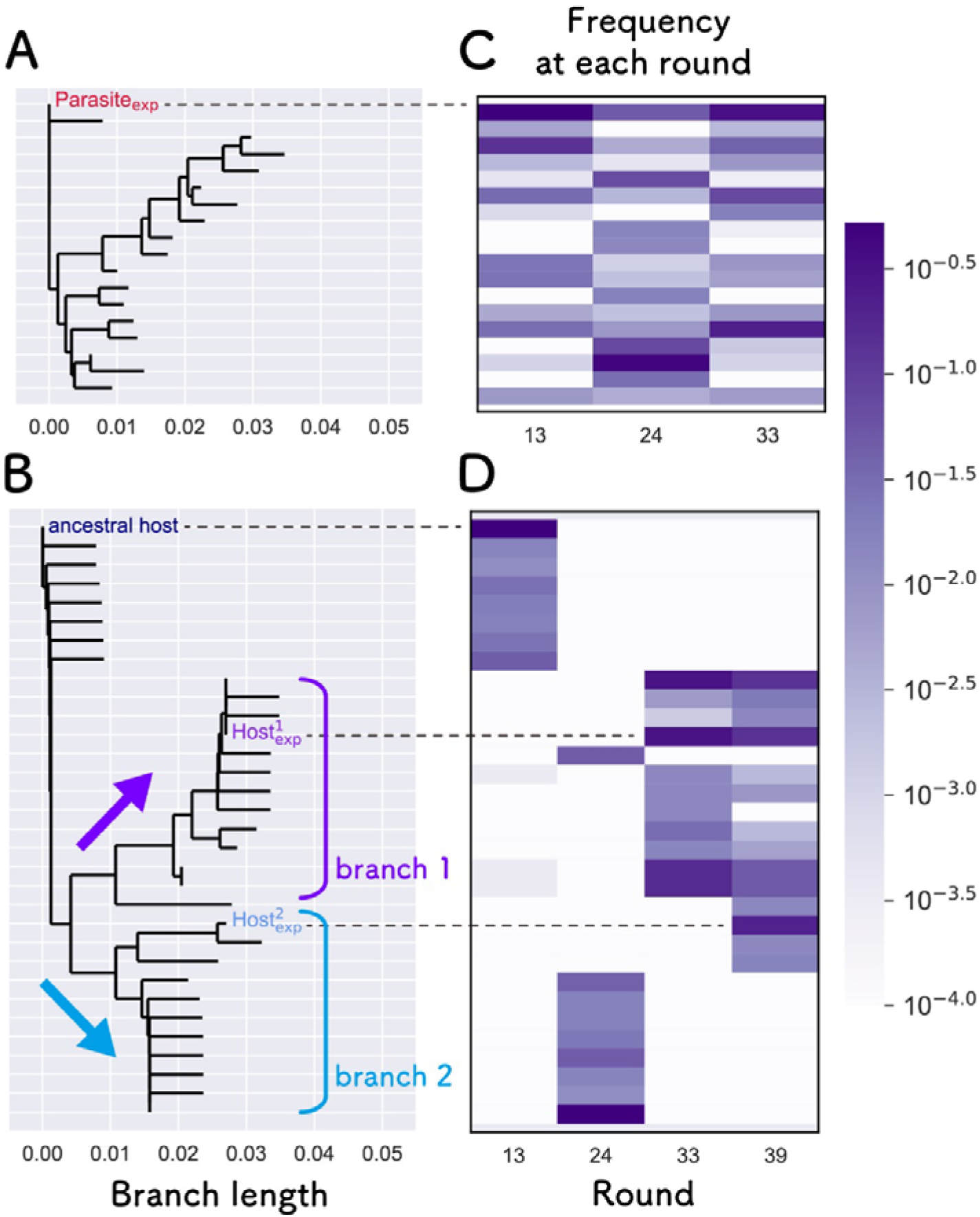
Phylogenetic analysis of the host and parasitic RNAs that appeared in the previous evolutionary experiment. Phylogenetic trees of the top eight parasitic (A) and host RNAs (B) that appeared in the early rounds of the previous evolutionary experiment (30). Phylogenetic trees were constructed using the neighbor-joining method with the Phylo.TreeConstruction module in the Biopython library and default parameters (46–48). The RNA frequencies at each round are shown as heat maps for the parasite (C) and host RNAs (D). Representative parasites and hosts used for the next biochemical experiments are indicated by “Parasite_exp_” and 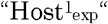 and 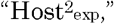 respectively. We could not obtain sequence data of the parasite at round 39 because the total concentration of the parasitic RNA was too low.

The phylogenetic tree of the host RNAs was divided into two major branches (Fig. 8B), consistent with the result of a previous study (30). The frequency of the host RNAs changed significantly in each round (Fig. 8D). At round 13, the sequences around the ancestral host dominated the population, and then, a part of the RNAs in branch 2 dominated the population at round 24. At round 33, the major RNA population changed to branch 1. At round 39, most of the RNAs in branch 1 remained as a major population, but some RNAs in branch 2 participated in the population as new dominant RNAs. From these results, we hypothesized that the dominant host RNA in branch 1 and a parasitic RNA form the HP network at round 33, which then changed to HHP network at round 39 by the addition of another host RNA in branch 2. To verify this hypothesis, we chose two representative hosts, one of the most frequent RNAs in branch 1 from round 33 to 39 (indicated as 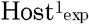) and the most common RNA in branch 2 at round 39 (indicated as 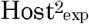).

### Parameter estimation of the representative RNAs

Next, we estimated the replication coefficients (*k_ij_*) of 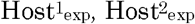, and Parasite_exp_. To measure the coefficients, we performed two-step reactions for all RNA combinations (Fig. 9A). In the first translation reaction, RNA replicase is translated from one of the host RNAs, and in the second replication reaction, the translated replicase was used for replication of the same host and/or another host or parasitic RNA. The results of replication are shown in Fig. 9B. The results indicated that parasite replication was asymmetric. When comparing the red bars, we found that the parasite was replicated when 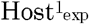 was used as RNA I (i.e., by 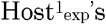 replicase), while it was barely replicated when 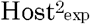 was used as RNA I (i.e., by 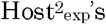 replicase), indicating that 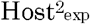 is more resistant to the parasite, consistent with the sustainable asymmetric case shown in Figs. 5C and 5D. To quantitatively compare the parameters, we estimated the replication coefficients from the replication results (Table 1). The replication coefficients of 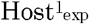 and the parasite are close to one of the sustainable conditions in the HP network (a light blue square in Fig. 3F) and on the edge of the sustainable conditions in the HHP network (a magenta square in Fig. 5D). These results indicate that the RNA species that appeared during the evolutionary experiment have properties that allow sustainable HP and HHP networks.

**Figure 9.**
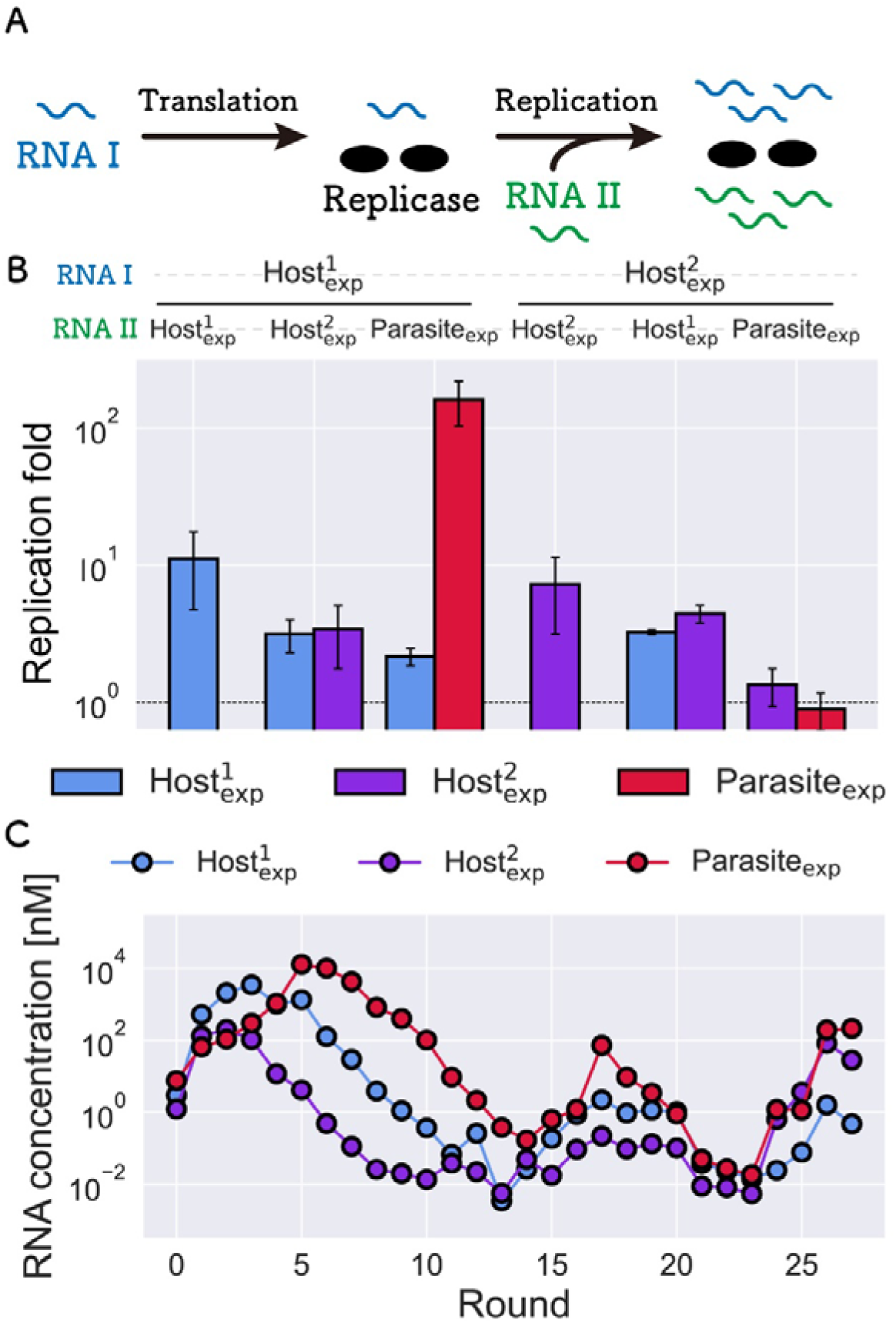
Biochemical analysis of the representative host and parasitic RNAs. (A) Experimental procedure for the estimation of replication coefficients. In the first translation reaction, RNA replicase was translated from one of the host RNAs (RNA I) for 2 h at 37°C, in which UTP was omitted to avoid RNA replication. In the second replication step, another host or parasitic RNA (RNA II), UTP, and an inhibitor of translation (30 μg/ml streptomycin) were added, and both RNAs I and II were replicated by the replicase for 1 h at 37°C. (B) RNA replication results. Experiments are independently performed three times. The error bars represent standard deviations. (C) Trajectory of RNA concentrations in the compartmentalized serial replication experiment of the three representative RNAs.

We further tested whether the representative host and parasite RNAs co-replicated sustainably using compartmentalized serial replication experiments. We mixed the three representative RNAs at an equivalent concentration (10 nM) in a cell-free translation solution and encapsulated them into water-in-oil droplets. The replication was repeated using the same serial replication procedure as in a previous study (30). All three RNAs were replicated until 27 rounds while maintaining detectable concentrations (Fig. 9C), supporting the notion that the selected host and parasite RNAs have the ability to form a sustainable HHP network.

## Discussion

In this study, we investigated the plausible complexification pathway of host–parasite replication networks using computer simulation and experiments. First, we examined the parameter space that allowed sustainable replication of all members in replication networks from two- (HH and HP) to three-member networks (HHH, HHP, and HPP). Sustainable parameter spaces are broader in HP and HHP networks for the range of the parameters we used, suggesting the plausibility of complexification from a single replicator to HP and then to HHP networks. We further confirmed that the dominant RNAs isolated from the previous evolutionary experiments had parameter sets that sustained HP and HHP networks, suggesting that the transition of replication network actually occurred during the evolutionary experiment. These results provide both theoretical and experimental evidence that the spontaneous development of a complex reaction network through Darwinian evolution is feasible within the parameter space that is achievable with RNA and proteins and that coevolution with parasitic replicators plays an important role in the complexification.

To date, the conditions required for the coexistence of multiple replicators have been studied using various theoretical models (15,34–37). The sustainable conditions found in this study are consistent with those of previous studies. For example, the sustainability of an HHP network that requires a parasite and asymmetric parasite resistance between the two hosts (Fig. 5C) is consistent with the idea that the parasites play a role as a “niche” to sustain different types of host species (15,36,37). In addition, the sustainability of HH networks that require larger cross-replication than self-replication (Fig. 3C) is similar to the cooperative relationship found in hypercyclic networks (17). This consistency with previous theoretical studies, however, does not diminish the importance of this study because the novelty of this study is not to provide a new concept for the coexistence theory but to reveal realistic pathways and parameters for the complexification in biochemical replicator systems. In this study, we found that the RNAs and the encoded replicase protein were able to have replication parameters that permit sustainable HP and HHP networks under compartmentalized conditions. We also found that experimentally obtained RNAs are on the edge of the sustainable parameter space (shown in the magenta square in Fig. 5D), which implies that a slight change in one of the parameters easily destroys the sustainability. The analysis of this study using an experimental model and relevant computer simulation revealed the realistic yet fragile nature of molecular replication networks.

We found that the HHH network can be sustainable in certain parameter spaces (Fig. 6B), suggesting that replication networks consisting of only host species can be another feasible complexification pathway. Such a replication network requires smaller self-replication and larger cross-replication values, and thus, it is similar to the hypercycles proposed by Eigen (17). However, such replication networks might be unlikely because they require a parasite-free environment. Parasitic replicators are reported to be inevitable in self-replicators with a certain level of complexity (39), as shown by the appearance of parasitic replicators soon after the initiation of replication in our translation-coupled RNA or DNA replication systems (29,39). Once parasitic replicators appear, the HH network changes to a more sustainable HHP network. Therefore, the pathway from HH to HHH networks is possible, but can be realized in limited replication systems where parasitic replicators rarely appear.

The importance of parasitic entities in diversifying host species through evolutionary arms race and its inevitability have been proposed in various organisms (38,40–42), digital organisms (43), and molecular replicators (15,30,31). The relatively broader parameter space that allows sustainable HHP network may imply that HHP network, in which the newly appeared host uses the parasite as a “niche,” is a reasonable consequence of coevolution between host and parasite. If this pathway continues, the network may further develop by acquiring a new parasite and then a new resistant host continuously (Fig. S5). The analysis of a more complex replication network that includes a larger number of replicators is a remaining challenge. Indeed, we recently reported that after 240 rounds of serial replication cycles, a five-member network consisted of three hosts and two parasites appeared, in which the host RNAs have asymmetric resistance to the two parasites (31). It is of utmost importance how many RNAs participate in a network, what determines the maximum number, and whether the different RNAs fuse to become a single molecule that encodes more information, which may lead to the origin of chromosome (21). The theoretical and experimental models used here provide a useful tool for answering these questions.

## Materials and Methods

### Simulation of compartmentalized replication through serial replication cycle

The replication was continued by repeating three steps: replication, selection, and fusion-division (Fig. 2A). In the replication step (Fig. 2B), hosts and parasitic replicators in each compartment replicate depending on their concentrations in each compartment according to the different equations described below. The replication reactions are described using the following logistic equations that take self- and non-self-replications among host and parasites into account:

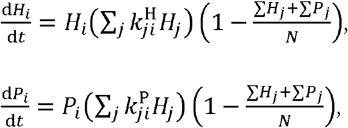

where *H* and *P* are the concentrations of the hosts and parasites, respectively. *k_ji_* is the coefficient of the reaction in which host *j* replicates the host or parasite. *N* is the carrying capacity of each compartment. In this equation, we assumed that the replication rate of each host or parasite depends on three factors: own concentration (*H_i_* or *P_i_*), the sum of the host concentration multiplied by its replication ability, (∑_*j*_ *k_ji_ H_j_*), which represents the total replication ability provided by host replicators in the compartment, and the effect of carrying capacity in the compartment 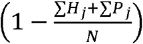. Compartments are assumed to be independent reactors, and there is no interaction between replicators in different compartments. The total number of compartments are fixed as C. The sizes of all the compartments were the same.

In the selection phase (Fig. 2C), a certain number (*C_S_*) of compartments was randomly selected, and empty compartments were supplied up to the fixed total number of compartments (*C*). The number of selected compartments (*C_S_*) is defined as *C_S_* = ⌊*C* × *S*⌋, where *S* (∈ [0, 1]) is the selection rate.

In the fusion-division phase (Fig. 2D), the following three steps are repeated *A* times (*A* is defined as “fusion-division frequency”). First, two compartments were randomly chosen from all compartments. Second, the concentrations of each replicator (i.e., hosts or parasites) in the two compartments were summed, and the number of replicators in the compartment was approximated from the concentrations. Third, the replicators were randomly redistributed into two new compartments, and their concentrations were calculated.

To search for parameter sets that allow sustainable replication, we conducted the replication-selection-fusion-division cycle for 100 rounds and counted each number of “sustained runs” out of 10 independent runs. We defined the “sustained run” as that where the number of all hosts or parasites in the network is greater than the number of compartments (i.e., all compartments contain all hosts or parasites on average) in the final round. In these simulations, all compartments were initially filled with equal numbers of all hosts and parasites, as much as the carrying capacity. The number of compartments (*C*) was 3,000, the selection rate (*S*) was 0.25, the fusion-division frequency (*A*) was 5,000, and the carrying capacity (*N*) was 100.

For the simulation of the evolutionary transition shown in Fig. 7, a mutation step was introduced just before the replication step, in which one of the compartments that contain hosts was randomly chosen and a new replicator (i.e., host or parasite) appeared in the compartments at a mutation rate 0.5 × *H_t_*, where *H_t_* is the total host concentration in the compartment. To reduce the computational cost, the total number of replicator species is restricted to less than three. That is, a new species can appear in the mutation step only when the total number of replicator species is one or two. A new replicator species that appears by mutation is a host or a parasite at the probabilities of 0.2 or 0.8, respectively. The coefficients of a new host are randomly chosen from 1.7, 1.8, 1.9, 2.0, 2.1, 2.2, 2.3, 2.4, 2.5, and 2.6. Coefficients of a new parasite were randomly chosen from 0.01, 0.1, 0.5, 1, 2, 3, 4, 5, 6, 7, 8, 9, and 10. The simulation was started from a single host that self-replicates with a coefficient 2.0 and continued for 3000 rounds of serial replication cycles. The number of compartments (*C*) was 3,000, the selection rate (*S*) was 0.25, the fusion-division frequency (*A*) was 5,000, and the carrying capacity (*N*) was 100.

### RNA preparation

The representative host and parasitic RNAs were prepared by in vitro transcription using each plasmid as previously described (44). The plasmids encoding each representative RNA (pUC_RK-Host-1, pUC_RK-Host-2, and pUC-RK-Parasite) were constructed in this study by introducing mutations into the plasmid pUC-N96 that encodes the original RNA by PCR with mutated primers. These mutations are listed in Table S1. All RNA sequences are shown in the supplemental text.

### Replication experiments and estimation of the parameters

The procedure was based on a previous study (31), which included two steps. First, a host RNA (30 nM, RNA I) was incubated at 37°C for 2 h in a cell-free translation system in which UTP was omitted to avoid RNA replication. The cell-free translation system is a reconstituted translation system of *Escherichia coli* (45). The composition was customized and reported in a previous study (29). Next, the initial reaction solution was diluted 3-fold in the cell-free translation system, which contains another RNA (10 nM, RNA II), 1.25 mM UTP, and 30 μg/mL streptomycin to inhibit further translation, followed by incubation at 37°C for 1 h. The mixtures were diluted 10,000 fold with 1 mM EDTA (pH 8.0) and each RNA concentration was measured by quantitative PCR after reverse transcription using PrimeScript One Step RT-PCR Kit (TaKaRa, Japan) with specific primers (Table S2). Reverse transcription was performed for 30 min at 42°C, followed by 10 s at 95°C. PCR was performed for 5 s at 95°C and 30 s at 60°C for 50 cycles.

To estimate replication coefficients, we first calculated the common logarithms of the increase ratios from 0 to 1 h as fold values, 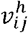, where the subscripts *i* and *j* represent RNA species used as RNA I and II, respectively, and the superscript *h* represents the measured RNA species (i or j). The fold values when the same host RNA was used for both RNA I and II were utilized as the self-replication coefficients (i.e., 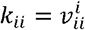). The fold values when different RNAs were used for RNA I and II were utilized as the nonself-replication coefficients after normalization to eliminate the competition effect between RNA I and II on replication according to the following equation:

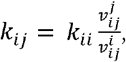

where we assumed that the ratio of the fold values of the competitive RNAs 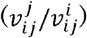 the same as the ratio of the self-replication coefficient (*k_ij_* /*k_ii_*).

### Compartmentalized serial replication experiment of the representative hosts and parasitic RNAs

The serial replication experiment shown in Fig. 9C was performed according to a previous study (29). Briefly, the initial reaction mixture contained 10 nM 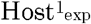, 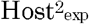, and Parasite_exp_ in the reconstituted translation system described above. The solution (10 μL) was dispersed in 1 mL of the saturated oil phase with a homogenizer (Polytron Pt-1300d; Kinematica) at 16,000 rpm for 1 min on ice and incubated for 5 h at 37°C. An aliquot (200 μL) of the droplets was diluted with 800 μL of the saturated oil phase, and a new solution of the reconstituted translation system was added. The solution was vigorously mixed with the homogenizer at 16,000 rpm for 1 min on ice and incubated for 5 h at 37°C. Thus, we repeated the serial replication cycle for 27 rounds. After incubation, the droplets were diluted 100-fold with 1 mM EDTA (pH 8.0) and each RNA concentration was measured by quantitative PCR after reverse transcription using PrimeScript One Step RT-PCR Kit (TaKaRa) with each specific primer (Table S2).

