## Supplemental Information for "Plausible pathway for a host–parasite molecular replication network to increase its complexity through Darwinian evolution"

**Supplemental Figures and Tables**

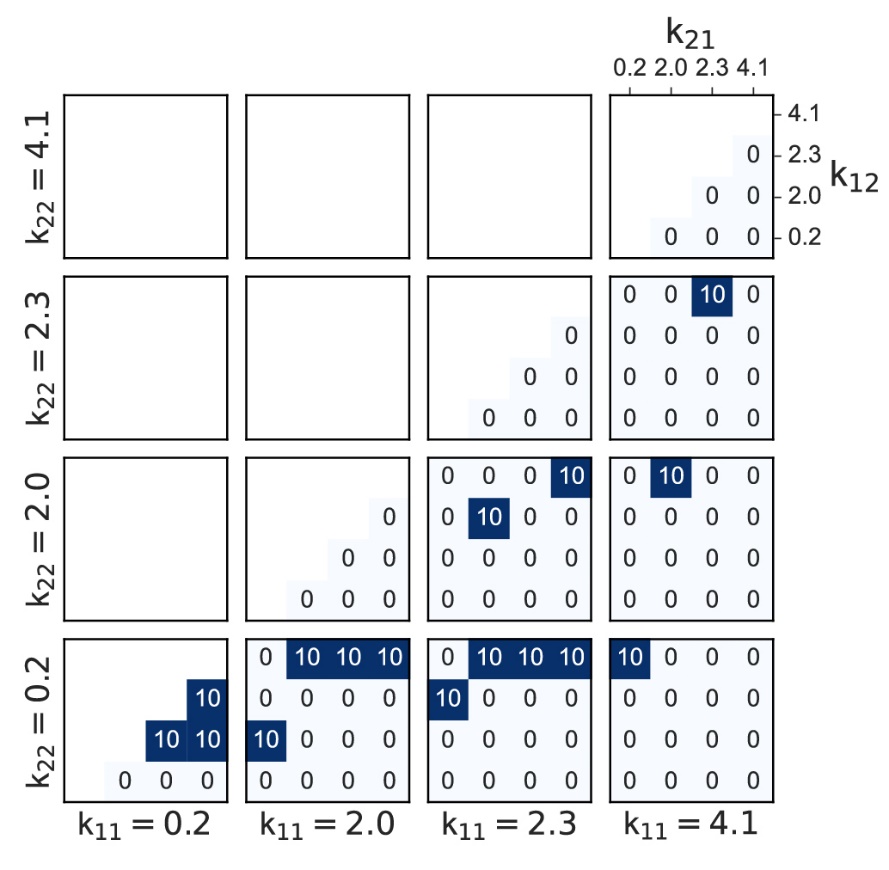

**Figure S1. Search for sustainable parameters in HH network with extreme parameters.** The simulation procedure was the same as that shown in Fig. 3B except for using smaller (0.2) and larger (4.1) parameter values.

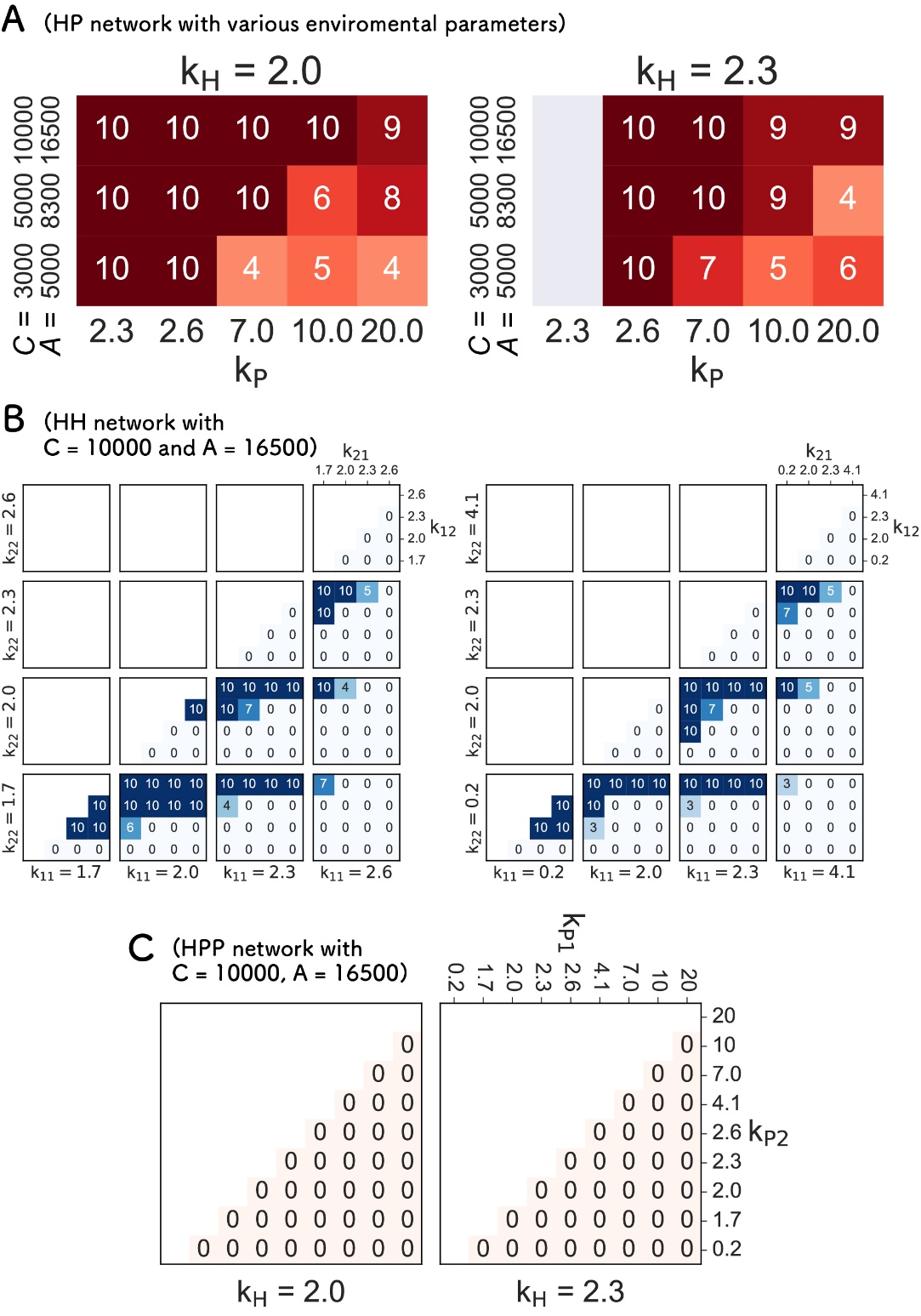

**Figure S2. Search for the parameters that allow sustainable HP, HH, and HPP networks with large numbers of compartments.**

The number of compartments and the frequency of fusion-division were increased to 10,000 and 16,500, respectively. The number of runs in which all three replicators (Hosts 1 and 2, and the parasite) were sustained for 100 rounds out of 10 independent runs are shown. (A) HP network. The replication coefficient for the host self-replication is fixed at 2.0 or 2.3. (B) HH network. (C) HPP network.

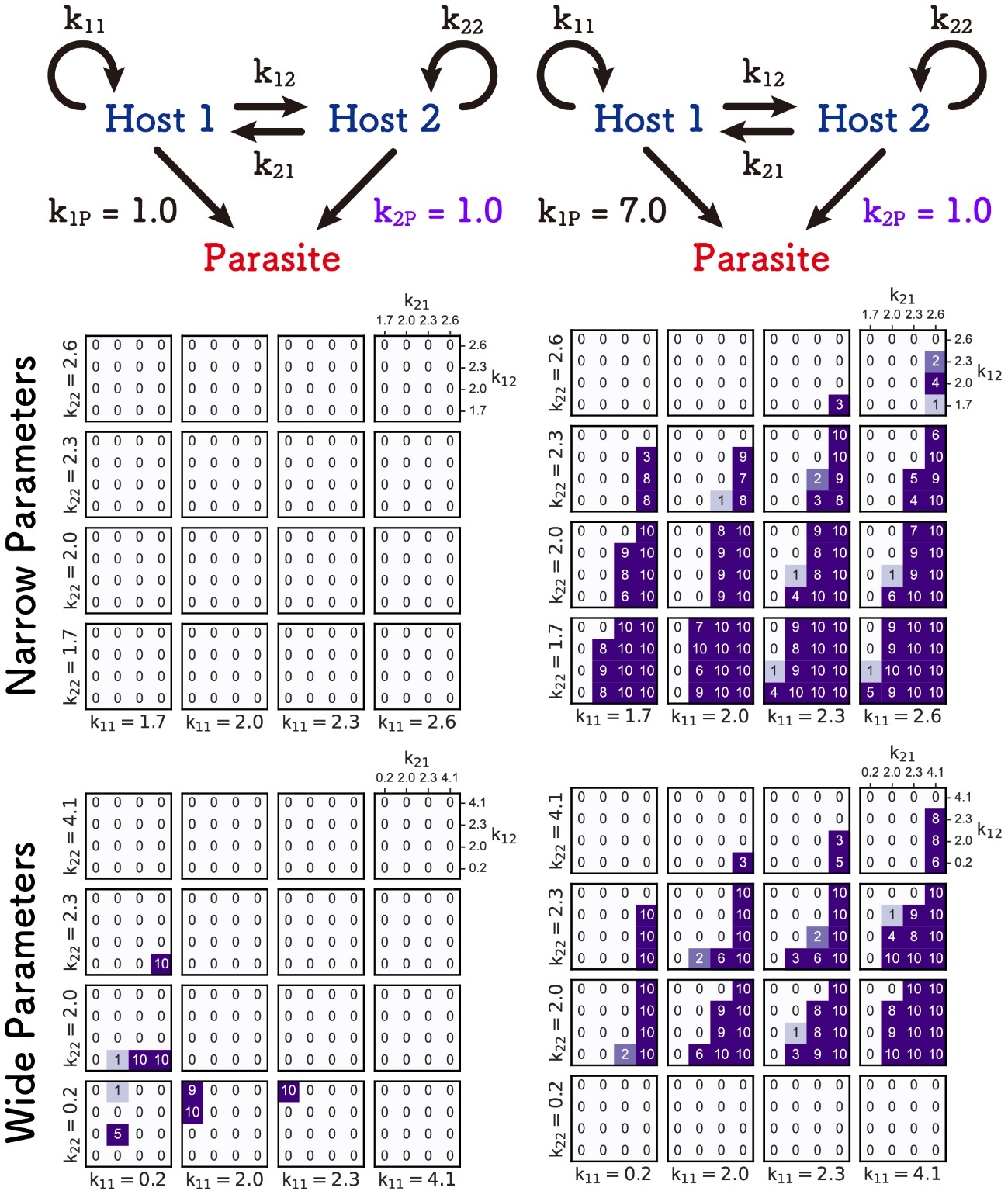

**Figure S3. Search for the parameters that allow sustainable asymmetrical HHP network with intermediate k_2P_ values.**

The simulations of the HHP network were conducted by the same method as Fig. 5 except for employing an intermediate k_2P_ values (1.0). The number of runs in which all three replicators (Hosts 1 and 2, and the parasite) were sustained for 100 rounds in 10 independent runs are shown.

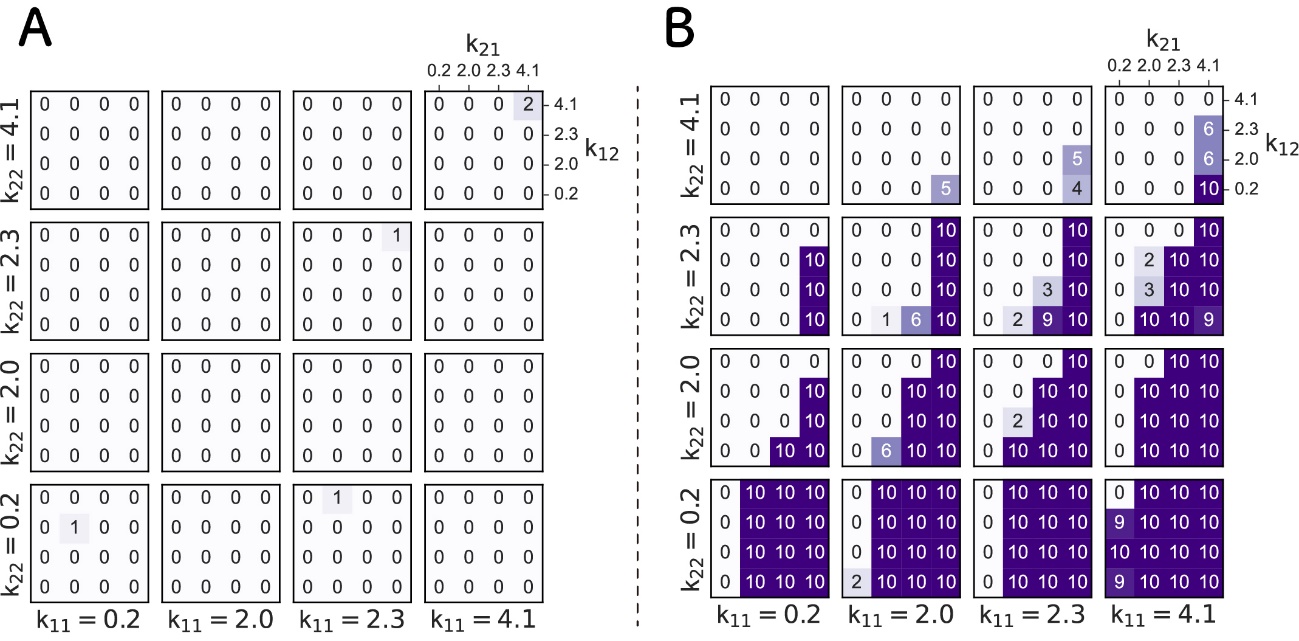

**Figure S4. Search for the parameters that allow sustainable HHP network with extreme parameters.**

The simulations of the HHP network were conducted in the symmetrical (A) or asymmetrical cases (B) by the same method as Fig. 5 except for employing extreme parameter values (0.2 an 4.1). The number of runs in which all three replicators (Hosts 1 and 2, and the parasite) were sustained for 100 rounds in 10 independent runs are shown.

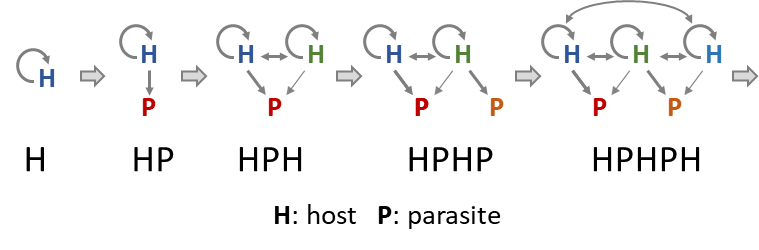

**Figure S5. A hypothetical parasite-mediated complexification pathway in replication networks**

**Table S1. Mutations in the representative host RNAs**

| Position* | Origin | Host 1 | Host 2 |
| --- | --- | --- | --- |
| 86 | C | C | U |
| 116 | A | G | A |
| 258 | C | C | U |
| 849 | A | G | A |
| 1566 | U | U | C |
| 1570 | U | G | U |
| 1603 | A | G | A |

*Positional numbers are based on the original host RNA. All sequences are shown in the Supplemental text.

**Table S2. Primer sequences**

| Target RNA | Direction | Sequence |
| --- | --- | --- |
| Host^1^_exp_ | Forward | TTGCTGCCTAAACAGCTGCG |
|  | Reverse | CGCTCTTGGTCCCTTGTATG |
| Host^2^_exp_ | Forward | TAGGGCCCATTCTGTGCACCT |
|  | Reverse | CGCTCTTGGTCCCTTGTATG |
| Parasite_exp_ | Forward | TACCGAAACGCACGAAGG |
|  | Reverse | CGTACGGGAGTTCGACCG |

**Supplemental text**

RNA sequences

> Original host

GGGAACCCCCCUUCGGGGGGUCACCUCGCGCAGCGGGCUACGCGAGGGAGCCACGCUGCGAAGCAGCGUGGCGGUUCUCGUGCGUCACCGAAACGCACGAAGGUCGCGCCUCUUCACGAGGCGUCACCUGGGAGAGCGCGAAAGCGCUAGCCCGUGCUCUAGCUCUAGAAGGUCUCGAGAUCUCCUCUAGAGAUAAUUUUGUUUAACUCUAAGAAGGAGAUAUACACAUGCCUAAGACAGCAUCUUCGCGUAACUCUCUCAGCGCACAAUUGCGCCGAGCCGCGAACACAAGAAUUGAGGCUGAAGGUAACCUCGCACUUUCCAUUGCCAACGAUUUACUGUUGGCCUAUGGUCAGUCGCCAUUUAACUCUGAGGCUGAGUGUAUUUCAUUCAGCCCGAGAUUCGACGGGACCCCGGAUGACUUUAGGAUAAAUUAUCUUAAAGCCGAGAUCAUGUCGAAGUAUGACGACUUCAGCCUAGGUAUUGAUACCGAAGCUGUUGCCUGGGAGAAGUUCCUGGCAGCAGAGGCUGAAUGUGCUUUAACGAACGCUCGUCUCUAUAGGCCUGACUACAGUGAGGAUUUCAAUUUCUCACUGGGCGAGUCAUGUAUACACAUGGCUCGUAGAAAAAUAGCCAAGCUAAUAGGAGAUGUUCCGUCCGUUGAGGAUAUGUUGCGUCACUGCCGAUUUUCUGGCGGUGCUACAACAACGAAUAACCGUUCGUACAGUCAUCCGUCCUUCAAGUUUGCACUUCCGCAAGCGUGUACGCCUCGGGCUUUGAAGUAUGUUUUAGCUCUCAGAGCUUCUACACAUUUCGAUAUCAGAAUUUCUGAUAUUAGCCCUUUUAAUAAAGCAGUUACCGUACCUAAGAACAGUAAGACAGAUCGUUGUAUUGCUAUCGAACCUGGUUGGAAUAUGUUUUUCCAACUGGGUAUCGGUGGCAUUCUACGCGAUCGGUUGCGUUGCUGGGGUAUCGAUCUGAAUGAUCAGACGAUAAAUCAGCGCCGCGCUCACGAAGGCUCCGUUACUAAUAACUUAGCAACGGUUGAUCUCUCAGCGGCAAGCGAUUCUAUGUCUCUUGCCCUCUGUGAGCUCUUAUUGCCCCGAGGCUGGUUUGAGGUUCUUAUGGACCUCAGAUCACCUAAGGGGCGAUUGCCUGACGGUAGUGUUGUUACCUACGAGAAGAUUUCUUCUAUGGGUAACGGUUACACAUUCGAGCUCGAGUCGCUUAUUUUUGCUUCUCUCGCUCGUUCCGUUUGUGAGAUACUGGACUUAGACUCGUCUGAGGUCACUGUUUACGGAGACGAUAUUAUUUUACCGUCCCGUGCAGUCCCUGCCCUCCGGGAAGUUUUUAAGUAUGUUGGUUUUACGACCAAUACUAAAAAGACUUUUUCCGAGGGGCCGUUCAGAGAGUCGUGCGGCAAGCACUACUAUUCUGGCGUAGAUGUUACUCCCUUUUACAUACGUCACCGUAUAGUGAGUCCUGCCGAUUUAAUACUGGUUUUGAAUAACCUAUAUCGGUGGGCCACUAUUGACGGCGUAUGGGAUCCUAGGGCCCAUUCUGUGUACCUCAAGUAUCGUAAGUUGCUGCCUAAACAGCUGCAACAUAAUACUAUACCUGACGGUUACGGUGAUGGUGCCCUCGUCGGAUCGGUCCUAAUCAAUCCUUUCGCGAAAAACCGCGGGUGGAUCCGGUACGUACCGGUGAUUACGGACCAUACAAGGGACCAAGAGCGCGCUGAGUUGGGGUCGUAUCUCUACGACCUCUUCUCGCGUUGUCUCUCGGAAAGUAACGAUGGGUUGCCUCUUAGGGGUCCAUCGGGUUGCGAUUUUGCUGAUCUAUUUGCCAUCGAUCAGCUUAUCUGUAGGAGUGAUCCUACGAAGAUAAGCAGGCCUACCGGUAAAUUCGAUAUACAGUACAUCGCGUGCUGUUGUUCGGGUUGUUGUUAGGCUUGCGGCCGCACUCGAGAGAUCUAGAGCAUCACGGUCGAACUCCCGUACGAGGUGCCCGCACCUCGUCCCCCCCUUCCGGGGGGGUCCCC

> Representative Host^1^exp

GGGAACCCCCCUUCGGGGGGUCACCUCGCGCAGCGGGCUACGCGAGGGAGCCACGCUGCGAAGCAGCGUGGCGGUUCUCGUGCGUCACCGAAACGCACGAAGGUCGCGCCUCUUCGCGAGGCGUCACCUGGGAGAGCGCGAAAGCGCUAGCCCGUGCUCUAGCUCUAGAAGGUCUCGAGAUCUCCUCUAGAGAUAAUUUUGUUUAACUCUAAGAAGGAGAUAUACACAUGCCUAAGACAGCAUCUUCGCGUAACUCUCUCAGCGCACAAUUGCGCCGAGCCGCGAACACAAGAAUUGAGGCUGAAGGUAACCUCGCACUUUCCAUUGCCAACGAUUUACUGUUGGCCUAUGGUCAGUCGCCAUUUAACUCUGAGGCUGAGUGUAUUUCAUUCAGCCCGAGAUUCGACGGGACCCCGGAUGACUUUAGGAUAAAUUAUCUUAAAGCCGAGAUCAUGUCGAAGUAUGACGACUUCAGCCUAGGUAUUGAUACCGAAGCUGUUGCCUGGGAGAAGUUCCUGGCAGCAGAGGCUGAAUGUGCUUUAACGAACGCUCGUCUCUAUAGGCCUGACUACAGUGAGGAUUUCAAUUUCUCACUGGGCGAGUCAUGUAUACACAUGGCUCGUAGAAAAAUAGCCAAGCUAAUAGGAGAUGUUCCGUCCGUUGAGGAUAUGUUGCGUCACUGCCGAUUUUCUGGCGGUGCUACAACAACGAAUAACCGUUCGUACAGUCAUCCGUCCUUCAAGUUUGCACUUCCGCAAGCGUGUACGCCUCGGGCUUUGAAGUAUGUUUUAGCUCUCAGAGCUUCUACACAUUUCGAUAUCAGAAUUUCUGAUAUUAGCCCUUUUAAUGAAGCAGUUACCGUACCUAAGAACAGUAAGACAGAUCGUUGUAUUGCUAUCGAACCUGGUUGGAAUAUGUUUUUCCAACUGGGUAUCGGUGGCAUUCUACGCGAUCGGUUGCGUUGCUGGGGUAUCGAUCUGAAUGAUCAGACGAUAAAUCAGCGCCGCGCUCACGAAGGCUCCGUUACUAAUAACUUAGCAACGGUUGAUCUCUCAGCGGCAAGCGAUUCUAUGUCUCUUGCCCUCUGUGAGCUCUUAUUGCCCCGAGGCUGGUUUGAGGUUCUUAUGGACCUCAGAUCACCUAAGGGGCGAUUGCCUGACGGUAGUGUUGUUACCUACGAGAAGAUUUCUUCUAUGGGUAACGGUUACACAUUCGAGCUCGAGUCGCUUAUUUUUGCUUCUCUCGCUCGUUCCGUUUGUGAGAUACUGGACUUAGACUCGUCUGAGGUCACUGUUUACGGAGACGAUAUUAUUUUACCGUCCCGUGCAGUCCCUGCCCUCCGGGAAGUUUUUAAGUAUGUUGGUUUUACGACCAAUACUAAAAAGACUUUUUCCGAGGGGCCGUUCAGAGAGUCGUGCGGCAAGCACUACUAUUCUGGCGUAGAUGUUACUCCCUUUUACAUACGUCACCGUAUAGUGAGUCCUGCCGAUUUAAUACUGGUUUUGAAUAACCUAUAUCGGUGGGCCACUAUUGACGGCGUAUGGGAUCCUAGGGCCCAUUCUGUGUACCGCAAGUAUCGUAAGUUGCUGCCUAAACAGCUGCGACAUAAUACUAUACCUGACGGUUACGGUGAUGGUGCCCUCGUCGGAUCGGUCCUAAUCAAUCCUUUCGCGAAAAACCGCGGGUGGAUCCGGUACGUACCGGUGAUUACGGACCAUACAAGGGACCAAGAGCGCGCUGAGUUGGGGUCGUAUCUCUACGACCUCUUCUCGCGUUGUCUCUCGGAAAGUAACGAUGGGUUGCCUCUUAGGGGUCCAUCGGGUUGCGAUUUUGCUGAUCUAUUUGCCAUCGAUCAGCUUAUCUGUAGGAGUGAUCCUACGAAGAUAAGCAGGCCUACCGGUAAAUUCGAUAUACAGUACAUCGCGUGCUGUUGUUCGGGUUGUUGUUAGGCUUGCGGCCGCACUCGAGAGAUCUAGAGCAUCACGGUCGAACUCCCGUACGAGGUGCCCGCACCUCGUCCCCCCCUUCCGGGGGGGUCCCC

> Representative Host^2^exp

GGGAACCCCCCUUCGGGGGGUCACCUCGCGCAGCGGGCUACGCGAGGGAGCCACGCUGCGAAGCAGCGUGGCGGUUCUCGUGCGUUACCGAAACGCACGAAGGUCGCGCCUCUUCACGAGGCGUCACCUGGGAGAGCGCGAAAGCGCUAGCCCGUGCUCUAGCUCUAGAAGGUCUCGAGAUCUCCUCUAGAGAUAAUUUUGUUUAACUCUAAGAAGGAGAUAUACACAUGCCUAAGACAGCAUCUUCGCGUAACUCUUUCAGCGCACAAUUGCGCCGAGCCGCGAACACAAGAAUUGAGGCUGAAGGUAACCUCGCACUUUCCAUUGCCAACGAUUUACUGUUGGCCUAUGGUCAGUCGCCAUUUAACUCUGAGGCUGAGUGUAUUUCAUUCAGCCCGAGAUUCGACGGGACCCCGGAUGACUUUAGGAUAAAUUAUCUUAAAGCCGAGAUCAUGUCGAAGUAUGACGACUUCAGCCUAGGUAUUGAUACCGAAGCUGUUGCCUGGGAGAAGUUCCUGGCAGCAGAGGCUGAAUGUGCUUUAACGAACGCUCGUCUCUAUAGGCCUGACUACAGUGAGGAUUUCAAUUUCUCACUGGGCGAGUCAUGUAUACACAUGGCUCGUAGAAAAAUAGCCAAGCUAAUAGGAGAUGUUCCGUCCGUUGAGGAUAUGUUGCGUCACUGCCGAUUUUCUGGCGGUGCUACAACAACGAAUAACCGUUCGUACAGUCAUCCGUCCUUCAAGUUUGCACUUCCGCAAGCGUGUACGCCUCGGGCUUUGAAGUAUGUUUUAGCUCUCAGAGCUUCUACACAUUUCGAUAUCAGAAUUUCUGAUAUUAGCCCUUUUAAUAAAGCAGUUACCGUACCUAAGAACAGUAAGACAGAUCGUUGUAUUGCUAUCGAACCUGGUUGGAAUAUGUUUUUCCAACUGGGUAUCGGUGGCAUUCUACGCGAUCGGUUGCGUUGCUGGGGUAUCGAUCUGAAUGAUCAGACGAUAAAUCAGCGCCGCGCUCACGAAGGCUCCGUUACUAAUAACUUAGCAACGGUUGAUCUCUCAGCGGCAAGCGAUUCUAUGUCUCUUGCCCUCUGUGAGCUCUUAUUGCCCCGAGGCUGGUUUGAGGUUCUUAUGGACCUCAGAUCACCUAAGGGGCGAUUGCCUGACGGUAGUGUUGUUACCUACGAGAAGAUUUCUUCUAUGGGUAACGGUUACACAUUCGAGCUCGAGUCGCUUAUUUUUGCUUCUCUCGCUCGUUCCGUUUGUGAGAUACUGGACUUAGACUCGUCUGAGGUCACUGUUUACGGAGACGAUAUUAUUUUACCGUCCCGUGCAGUCCCUGCCCUCCGGGAAGUUUUUAAGUAUGUUGGUUUUACGACCAAUACUAAAAAGACUUUUUCCGAGGGGCCGUUCAGAGAGUCGUGCGGCAAGCACUACUAUUCUGGCGUAGAUGUUACUCCCUUUUACAUACGUCACCGUAUAGUGAGUCCUGCCGAUUUAAUACUGGUUUUGAAUAACCUAUAUCGGUGGGCCACUAUUGACGGCGUAUGGGAUCCUAGGGCCCAUUCUGUGCACCUCAAGUAUCGUAAGUUGCUGCCUAAACAGCUGCAACAUAAUACUAUACCUGACGGUUACGGUGAUGGUGCCCUCGUCGGAUCGGUCCUAAUCAAUCCUUUCGCGAAAAACCGCGGGUGGAUCCGGUACGUACCGGUGAUUACGGACCAUACAAGGGACCAAGAGCGCGCUGAGUUGGGGUCGUAUCUCUACGACCUCUUCUCGCGUUGUCUCUCGGAAAGUAACGAUGGGUUGCCUCUUAGGGGUCCAUCGGGUUGCGAUUUUGCUGAUCUAUUUGCCAUCGAUCAGCUUAUCUGUAGGAGUGAUCCUACGAAGAUAAGCAGGCCUACCGGUAAAUUCGAUAUACAGUACAUCGCGUGCUGUUGUUCGGGUUGUUGUUAGGCUUGCGGCCGCACUCGAGAGAUCUAGAGCAUCACGGUCGAACUCCCGUACGAGGUGCCCGCACCUCGUCCCCCCCUUCCGGGGGGGUCCCC

> Representative Parasite_exp_

GGGAACCCCCCUUCGGGGGGUCACCUCGCGCAGCGGGCUGCGCGAAGGAGCCACGCUGCGAAGCAGUGUGGCGGUUCUCGUGCGUUACCGAAACGCACGAAGGUCGCGCCUCUUCACGAGGCGUCACCUGGGAGAGCGCGAAAGCGCUAGCCCGUGAUUCGUCACGGUCGAACUCCCGUACGAGGUGCCCGCACCUCGUCCCCCCCUUCCGGGGGGGUCCCC
